# Susceptibility of raccoon dogs for experimental SARS-CoV-2 infection

**DOI:** 10.1101/2020.08.19.256800

**Authors:** Conrad M. Freuling, Angele Breithaupt, Thomas Müller, Julia Sehl, Anne Balkema-Buschmann, Melanie Rissmann, Antonia Klein, Claudia Wylezich, Dirk Höper, Kerstin Wernike, Andrea Aebischer, Donata Hoffmann, Virginia Friedrichs, Anca Dorhoi, Martin H. Groschup, Martin Beer, Thomas C. Mettenleiter

**Author notes:** Authors contributed equally to this work (Conrad M. Freuling, Angele Breithaupt). Corresponding authors: Address for correspondence: Martin Beer, Institute of Diagnostic Virology, Friedrich-Loeffler-Institut, Südufer 10, 17493 Greifswald-Insel Riems, Germany, and Thomas C. Mettenleiter, President, Friedrich-Loeffler-Institut, Südufer 10, 17493 Greifswald-Insel Riems, Germany.

## Abstract

Severe acute respiratory syndrome coronavirus 2 (SARS-CoV-2) emerged in China at the end of 2019, and became pandemic. The zoonotic virus most likely originated from bats, but definite intermediate hosts have not yet been identified. Raccoon dogs (*Nyctereutes procyonoides*) are kept for fur production, in particular in China, and were suspected as potential intermediate host for both SARS-CoV6 and SARS-CoV2. Here we demonstrate susceptibility of raccoon dogs for SARS-CoV-2 infection after intranasal inoculation and transmission to direct contact animals. Rapid, high level virus shedding, in combination with minor clinical signs and pathohistological changes, seroconversion and absence of viral adaptation highlight the role of raccoon dogs as a potential intermediate host. The results are highly relevant for control strategies and emphasize the risk that raccoon dogs may represent a potential SARS-CoV-2 reservoir. Our results support the establishment of adequate surveillance and risk mitigation strategies for kept and wild raccoon dogs.

**Article Summary Line:** Raccoon dogs are susceptible to and efficiently transmit SARS-CoV2 and may serve as intermediate host

## Text

Coronaviruses can infect a wide variety of animals, and are responsible for human diseases including severe acute respiratory syndromes (SARS). Both SARS coronavirus (SARS-CoV) *(1, 2)* and Middle East Respiratory Syndrome coronavirus (MERS-CoV) *(3, 4)*, are ß-coronaviruses and presumably originate from bats *(5)*. They likely adapted to other reservoir hosts like Asian palm civets (*Paradoxurus hermaphroditus*) *(6)* and dromedary camels (*Camelus dromedarius*) *(7)*. Natural SARS-CoV infections were also detected in raccoon dogs (*Nyctereutes procyonoides*) which, among other candidate species, have been discussed as a possible intermediate host for the first SARS-pandemic of 2002/2003 *(8)*.

The current SARS-CoV-2 pandemic started from Wuhan, China, at the end of 2019. Close relatives to SARS-CoV-2 were found in bats *(9)*, and pangolins (*Pholidota spp*) *(10, 11)*. Furthermore, spill-over infections to different carnivores (dogs, cats, lions, tigers and minks) were reported *(12, 13)*. However, whether the pandemic started by a direct transmission of the SARS-CoV-2 ancestor from bats to humans or via an intermediate mammalian host with further adaptation, is still under debate *(14)*. For both, SARS-CoV and MERS-CoV, intermediate hosts played a crucial role in transmission to humans. However, no definite intermediate host for SARS-CoV-2 has been identified up to now *(14)*, but animal species like pangolins, palm civets, or raccoon dogs are discussed *(15–17)*. Although the pandemic is driven by direct human-to-human transmission, case studies demonstrate that anthropo-zoonotic infections occurred by contact of infected humans with companion animals and farmed minks kept for fur production in the Netherlands, Denmark and Spain *(13, 18, 19)*. There is also evidence for zoo-anthroponotic infection of humans *(13)*.

Natural infections of raccoon dogs with SARS-CoV were reported *(8)*, indicating a potential role in the previous SARS-CoV epidemic. In fact, 14.14 million captive raccoon dogs held in China for fur production *(20)* represent 99% of the global share (Figure. S3A). However, experimental infections of these animals with SARS-CoV or SARS-CoV-2 under controlled conditions and serologic surveillance of kept or wild raccoon dogs have not been documented.

Since SARS-CoV and SARS-CoV-2 employ the same receptor molecule ACE2 for contact with the receptor-binding-domain (RBD) of the spike (S) protein *(21)*, a similar range of susceptible host species can be assumed. Molecular studies indicate that the ACE2 proteins of raccoon dogs can also serve as an efficient receptor for SARS-CoV *(2)* and SARS-CoV-2 *(15)*.

Following a previously established study design *(3)* (Figure. 1A), we tested the susceptibility of raccoon dogs to SARS-CoV-2. Nine animals were challenged by intranasal inoculation of 10^5^ TCID50 SARS-CoV-2 2019_nCoV Muc-IMB-1, and 3 additional animals were introduced at one day post-infection (dpi) to evaluate direct viral transmission.

**Figure 1.**
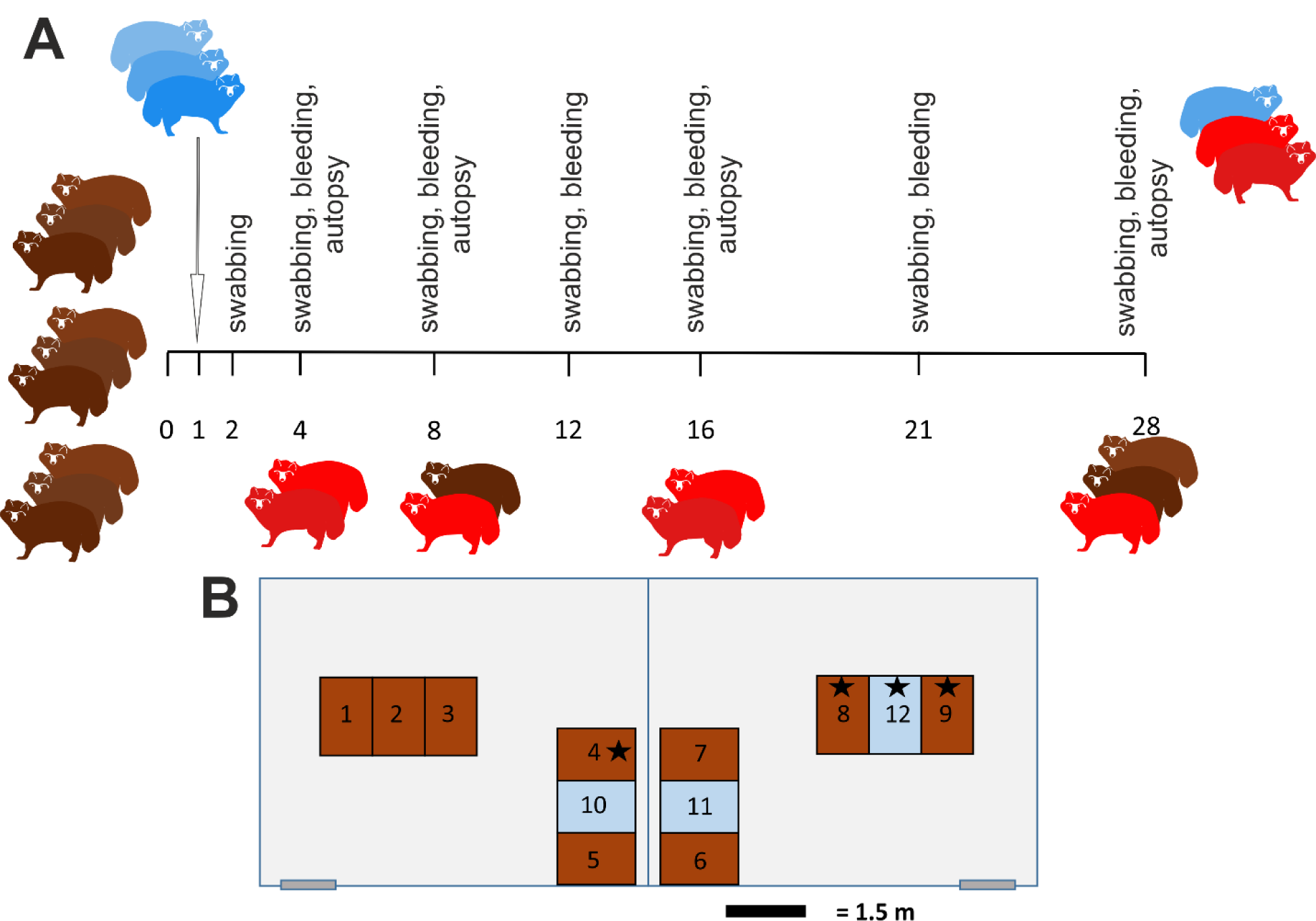
Study design (A) Outline of the in vivo experiments with an observation period of 28 days. Animals (n=9) were inoculated intranasally with 10^5^ TCID_50_/ml and three naïve direct contacts were added 1 dpi. On day 4 (animals #1, #2), day 8 (#3, #4), and 12 (#5, #6) two raccoon dogs each were sacrificed and subjected to autopsy. All remaining animals were euthanized on day 28 pi. Animals that became infected are highlighted in red. (B) Arrangement of the individual cages for the raccoon dogs in two separate rooms of the BSL 3 facility at the Friedrich-Loeffler-Institut. Inoculated animals (brown), contact animals (blue) and animals that remained uninfected 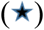 are indicated.

## Methods

### Study design

Fourteen adult, male (n=4) and female (n=10) raccoon dogs originating from a commercial farm were used. All animals were tested negative by RT-qPCR and antibody tests (ELISA, indirect immunofluorescence assay iIFAT, virus neutralization test VNT) for SARS-CoV-2 prior to the experiment. All raccoon dogs had been vaccinated against distemper, adeno-and parvovirus (Eurican^®^ SHP, Merial, France). Animals were kept in individual stainless-steel cages (1.5m × 0.95m × 2.0m) in four separate segments at 20°C room temperature, 60-80% humidity and a 12hr/12hr (35% dimming during night modus) lighting control within a fan forced draught ventilation equipped BSL3** animal facility at the Friedrich-Loeffler-Institut (FLI). Water was offered ad libitum. Animals were fed daily with 400 gr commercially produced feed for farmed foxes and raccoon dogs (Schirmer und Partner GmbH Co KG, Germany; Michael Hassel GmbH, Langenargen, Germany). The diet was supplemented with vitamins, minerals and items like one-day old chickens as described before *(24)*. The general health status of all animals, feed uptake and defecation were recorded daily. The body weight and temperature of all animals were measured prior to inoculation and at days 2, 4, 8, 12, 16, 21, and 28 pi.

The outline of the experiments with an observation period of 28 days is depicted in Figure 1. Nine raccoon dogs (3 males, 6 females) were infected intranasally with 10^5^ TCID_50_ SARS-CoV-2 2019_nCoV Muc-IMB-1. The inoculum of 2×1ml was administered to both nostrils using a pipette. To test viral transmission by direct contact, three naïve sentinel animals (all female) were added 24 hours post inoculation. Nasal, oropharyngeal and rectal swabs were taken at 2, 4, 8, 12, 16, 21 and 28 days post infection (dpi), blood was taken at 4, 8, 12, 16, 21 and 28 dpi. Two animals each were sacrificed at day 0 (control #1, #2) day 4 (animals #1, #2), day 8 (animals #3, #4) and day 12 pi (animals #5, #6). The remaining inoculated animals (animals #7-9) and the contacts (animals #10-12) were euthanized 28 dpi. All animals were subjected to autopsy for macroscopic evaluation and tissue sampling.

The animal experiments were evaluated and approved by the ethics committee of the State Office of Agriculture, Food Safety, and Fishery in Mecklenburg – Western Pomerania (LALLF M-V: LVL MV/TSD/7221.3-2-010/18-12).

### Virus and cells

The virus was propagated once in Vero E6 cells in a mixture of equal volumes of Eagle MEM (Hanks’ balanced salts solution) and Eagle MEM (Earle’s balanced salts solution) supplemented with 2mM L-Glutamine, nonessential amino acids, adjusted to 850 mg/L, NaHCO3, 120 mg/L sodium pyruvate, 10% fetal bovine serum (FBS), pH 7.2. No contaminants were detected within the virus stock preparation by metagenomic analysis employing previously published high throughput sequencing procedures Ion Torrent S5XL instrument *(25, 26)* and the sequence identity of the passaged virus (study accession number: PRJEB39640) was confirmed. The virus was harvested after 72 h, titrated on Vero E6 cells and stored at −80°C until further use.

### RNA extraction and detection of SARS-CoV-2

Total RNA from nasal, oropharyngeal and rectal swab samples, fecal samples as well as from tissues taken at autopsy were extracted using the NucleoMagVet kit (Macherey&Nagel, Düren, Germany) according to manufacturer’s instructions. Tissues were homogenized in 1 ml cell culture medium (see above) and a 5 mm steel bead in a TissueLyser (Qiagen, Hilden, Germany). Fecal samples were vortexed in sterile NaCl and the supernatant was sterile filtered (22µm) after centrifugation. Swab samples were transferred into 1 ml of serum-free tissue culture media and further processed after 30 min shaking. SARS-CoV-2 RNA was detected by an E-gene based RT-qPCR *(27)* using the AgPath-ID-One-Step RT-PCR kit (Thermo Fisher Scientific, Waltham, Massachusetts, USA) in a volume of 12.5 µl including 1 µl of ß-Actin-mix2-HEX as internal control and 2.5 µl of extracted RNA. The reaction was performed as described before *(23)*. Nasal swab samples from raccoon dog #2 (2 dpi) and from contact animal #10 (8 dpi) were subjected to high-throughput sequencing and compared to the inoculum (study accession number: PRJEB39640) by employing previously published high throughput sequencing procedures using Ion Torrent S5XL instrument *(25, 26)*.

### Detection of SARS-CoV-2 reactive antibodies

Serum samples collected throughout the study were tested for the presence of SARS-CoV-2 reactive antibodies by indirect immunofluorescence assay (iIFA), virus neutralization test (VNT) as described before *(23)*. For ELISA, medium-binding ELISA plates (Greiner Bio-One GmbH, Germany) were coated with SARS-CoV-2 RBD (RBD-SD1 domain, amino acids 319 – 519 of the SARS-CoV2 Spike ectodomain, for details see Appendix). The sera were diluted 1:100 in TBST and incubated on the coated and uncoated wells for 1h at room temperature followed by three washes using TBST. The saliva samples were used undiluted. Reactivity was shown by adding a multi species conjugate (SBVMILK; IDvet, France) diluted 1:80 (serum) or 1:10 (saliva). After an incubation period of 1 h at room temperature, the plates were washed again and Tetramethylbenzidine (TMB) substrate (IDEXX, Switzerland) was added. The ELISA readings were taken at a wavelength of 450 nm on a Tecan Spectra Mini instrument (Tecan Group Ltd, Switzerland). The measurements were normalized to the respective samples tested on wells treated only with the coating buffer.

For comparison, sera were also tested in a newly developed commercial SARS-CoV-2 sVNT *(28)*. Briefly, 1:10 serum dilutions were incubated for 30 min at 37°C with HRP-coupled RBD before transferring the samples to the capture plate pre-coated with the human ACE2 protein. After 15 min incubation at 37°C, plates were washed four times. TMB substrate was added and the plate was incubated at room temperature for 15 min before stopping the reaction and reading the optical density (OD) at 450 nm. Percent inhibition was calculated as (1-OD sample / OD negative control) × 100.

### Identification of SARS-CoV-2 RBD-specific immunoglobulins

SARS-CoV-2 specific immunoglobulins (Ig) were comparatively investigated in sera and saliva of raccoon dogs by ELISA using exactly the same SARS-CoV-2 RBD-SD1 antigen-coated plates, serum dilutions, washing and dilution buffers, TMB substrate, incubation periods and ELISA-Reader as described above. After the incubation of the sera or saliva and the following washing, dog-specific, horseradish-peroxidase (HRP) labelled Ig antibodies (goat-α-dog-IgA 1:1,000 for saliva and 1:5,000 for serum; goat-α-dog-IgM1 1:15,000; goat-α-dog-IgG, goat-α-dog-IgG1, goat-α-dog-IgG2 all 1:20,000; Bethyl Laboratories INC) were added and incubated for 1h at RT. Antibodies were diluted in TBST.

### Virus titration

Virus titer used for infection experiments was confirmed by titration on Vero E6 cells (Biobank Friedrich-Loeffler-Institut, catalogue N° 0929) and evaluation of CPE after 5 days. RT-qPCR positive swabs and tissue samples were titrated on Vero E6 cells as well

### Autopsy, histopathology, immunohistochemistry

Full autopsy was performed on all animals under BSL3 conditions. A broad spectrum of tissues was collected and fixed in 10% neutral-buffered formalin and trimmed for paraffin embedding, including the upper and lower respiratory tract, the gastro-intestinal tract, the urinary tract, brain, and main parenchyma (see appendix for details). Tissues were embedded in paraffin, and 3 μm sections were stained with hematoxylin and eosin for light microscopical examination. For SARS-CoV-2 antigen detection was performed as described *(25)*. Evaluation and interpretation of pathology data were performed by a board-certified pathologist (DiplECVP). The severity of lesions and the distribution of SARS-CoV-2 antigen was recorded on an ordinal scoring scale with scores 0 = no lesion/antigen, 1 = rare, affected cells/tissue <5% per slide; 2 = multifocal, 6-40 % affected; 3 = coalescing, 41-80% affected; 4 = diffuse, >80%affected.

### Statistical information

All data were analyzed and visualized using GraphPad Prism Version 7.0 (GraphPad Software, San Diego, CA, USA). No statistical methods were used.

## Results

Inoculation led to productive infection in six out of nine exposed animals. Based on the lack of viral RNA detection throughout the observation period of 28 days, we concluded that infection of animals #4, #8 and #9 failed (Figure 1, panel B). While several animals showed reduced overall activity at 4 dpi (animal #4, #5, #10), none of the exposed and contact animals showed any obvious clinical sign of infection until the termination of the experiment. In particular, neither increase in body temperature nor weight loss were observed.

Next, we examined the presence of viral RNA and infectious virus in nasal, oropharyngeal and rectal swab samples as well as in feces by quantitative reverse transcription PCR (RT–qPCR) and titration on Vero E6 cells. Raccoon dogs started to shed virus already at 2 dpi in nasal and oropharyngeal swabs (Figure 2, panels A, B). While infectious virus was isolated from individual animals up to 4 dpi (Figure. 2B), viral RNA was present in nasal swabs up to 16 dpi (animal #7, Figure 2, panel C). Viral genome loads were highest in nasal swabs (mean genome copies Log10/ml: 3.2, min: 1.0, max: 6.45), followed by oropharyngeal swabs (2.9; 0.54-4.39) and rectal swabs (0.71; 0.31-1.38, Figure 2, panel A). Virus titrations revealed the same trend, with the highest viral titres of up to 4.125 Log_10_ TCID_50_/ml in nasal swabs at 2 dpi. Infectious virus was never isolated from rectal swabs. In general, virus isolation failed above quantification cycle (Cq)-values of around 27 (Appendix Figure 1).

**Figure 2.**
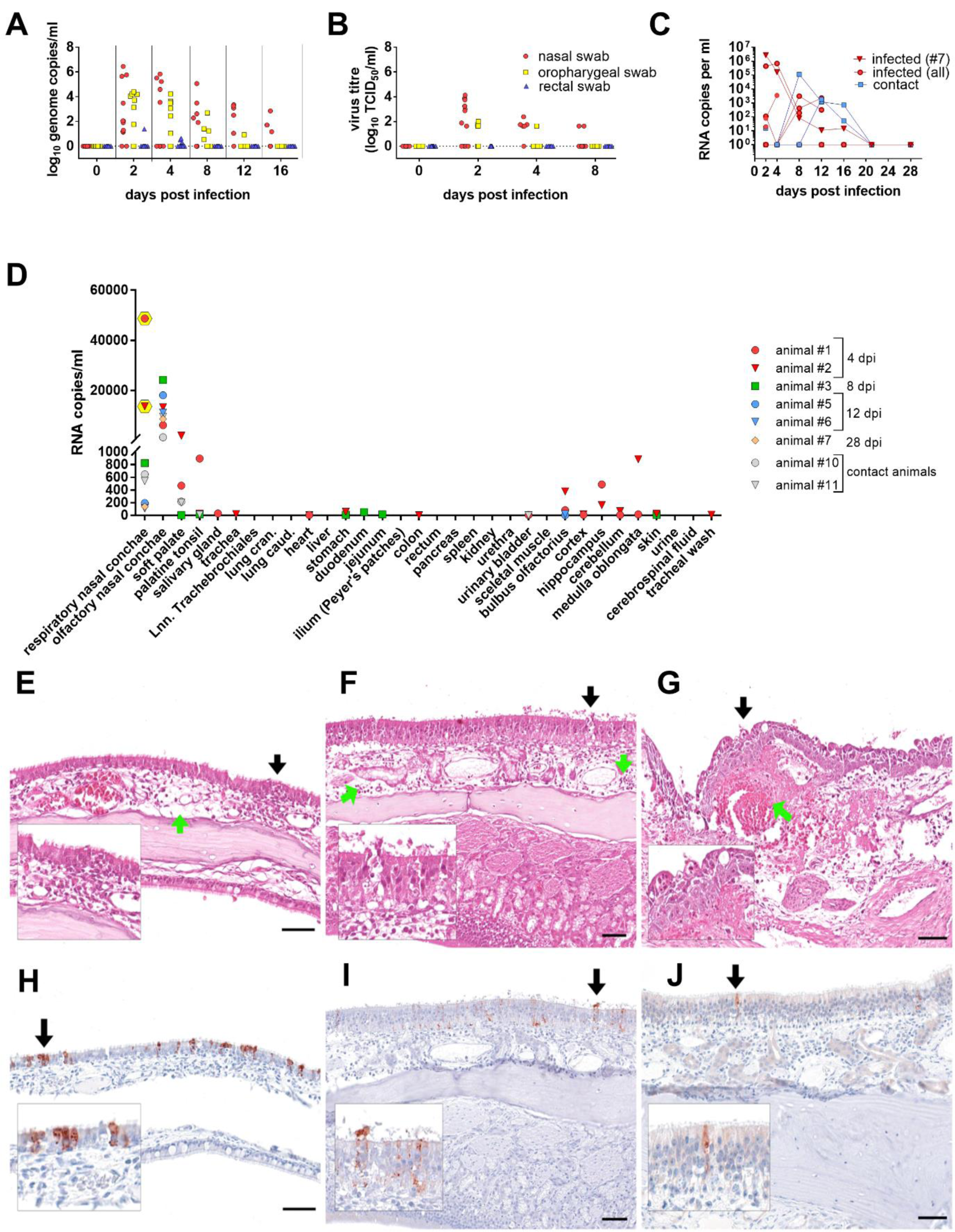
Virus detection in swab and tissue samples (A) SARS-CoV-2 viral genome loads in swab samples over time, and (B) virus titres using isolation on Vero E6 cells. Two replicates per sample were analysed and both results are shown. (C) Individual viral loads of nasal swabs taken from infected and contact animals. (D) Viral genome loads in organs, infectious virus was isolated only from nasal conchae at 4 dpi (yellow hexagons) from animal #1 (2.86 Log_10_ TCID50/ml) and animal #2 (1.63 Log_10_ TCID_50_/ml. (E) Rhinitis, respiratory region, with mucosal edema (green arrow) and epithelial degeneration with inflammation (black arrow, see also inlay) at 4 dpi. (F) Rhinitis, olfactory region, with mucosal edema and inflammation (green arrows) and epithelial necrosis and loss with minimal intraluminal debris (black arrow, see also inlay), 4 dpi. (G) Rhinitis, olfactory region, focal coagulative necrosis and hemorrhage (green arrow) and epithelial necrosis with early re-epithelisation (black arrow, see also inlay), 8 dpi. (H) Intralesional viral antigen oligofocal in the respiratory epithelium, 4 dpi, (I) Intralesional antigen labelling oligofocal in the olfactory epithelium, 4 dpi. (J) Single antigen-positive olfactory cells, 8 dpi. (E-G) Histopathology, hematoxylin & eosin stain, (H-J) immunohistochemistry, ABC method, AEC chromogen (red-brown), Mayer’s hematoxylin counter stain (blue). All bars = 50 μm

Virus was transmitted to two of three contact animals (#10, #11) (Figure 1, panel B, Figure 2, panel C). One contact raccoon dog (#12) remained negative due to the fact that both cage neighbors (#8, #9, Figure 1, panel B), did not shed virus after inoculation. In contact animals, viral RNA indicative of infection was first detected at 8 dpi (7 days post contact (dpc), #10). As in the inoculated animals, excretion in contact animals was mainly via nasal secretions and lasted until 16 dpi (15 dpc) and virus isolation yielded viral titers of 1.625 Log_10_ TCID_50_/ml in nasal swabs of one contact animal (#10) at 8 dpi (7 dpc).

Tissues and body fluids of euthanized animals were tested for the presence of SARS-CoV-2 RNA and replicating virus at day 4, 8, 12, and 28 pi (Figure 2, panel D). Highest viral genome loads of up to 4.87 Log10 genome copies per ml were observed in samples from the oro-nasal cavity, whereas only minute amounts were sporadically identified in other organs. The caudal, olfactory region of the oro-nasal cavity in general yielded higher viral genome loads than the cranial, respiratory region. Infectious virus could be cultivated from the nasal conchae of animals #1 (2.86 Log_10_ TCID_50_/ml) and #2 (1.63 Log_10_ TCID_50_/ml). Of note, none of the lung samples was positive for viral RNA, nor did any of the animals demonstrate viremia. Both animals investigated at 4 dpi had viral RNA in samples of the CNS with low genome loads (max 2.95 Log10 genome copies/ml), but cerebrospinal fluid was negative in all tested animals.

At autopsy, no gross lesions were recorded that could be assigned to the SARS-CoV-2-infection. However, histopathology identified mild rhinitis at 4, 8, and 12 dpi (animals #1-3, #5, #6). The olfactory, caudal region of the nasal cavity was more consistently affected compared to the respiratory, cranial region and included degeneration, necrosis and loss of the respiratory and olfactory epithelium, presence of intraluminal cellular debris, degeneration and necrosis of the submucosal glands, mucosal edema, endothelial swelling and acute, submucosal hemorrhage (Figure 2, panels E-G). At 4 dpi, mainly neutrophils with fewer macrophages and lymphocytes were found, later lesions showed predominantly lymphocytes and fewer neutrophils and macrophages. Mucosal coagulative necrosis with early re-epithelization and granulation tissue formation was present in one case (Figure 2, panel G, 8 dpi). At 28 dpi, one infected (#7) and one contact animal (#10) showed lesions indicative for previous viral replication sites in the nasal conchae. Viral RNA was still present, but no viral antigen was found (Appendix Figure 2).

Immunohistochemistry verified the presence of viral antigen only in the nasal conchae. Lesion associated antigen was found to be oligofocal at day 4 (animal #1, #2) in the respiratory and olfactory epithelium and to a lesser extent at day 8 (animal #3) only in the olfactory epithelium (Figure 2, panels H-J). No viral antigen could be found at 12 dpi and 28 dpi, and neither histopathologic lesions nor viral antigen were detected in animal #4 (8 dpi) in the nasal cavity. All other tissues tested negative for SARS-CoV-2 antigen.

SARS-CoV-2-specific antibodies were detected in all infected animals at 8 dpi as shown by ELISA (Figure 3, panels A-G) and iIFAT (> 1:64, Table 1). Titers increased up to 1:1024, detected at 28 dpi via iIFAT (animal #7). Neutralizing antibodies (VNA) were observed in two of the inoculated animals (#6, #7) as early as 8 dpi (#6, 1:5.04, Tab. 1). The highest VNA titer was 1:12.7 (#6, day 12; #7 day, 28). Interestingly, animal #7 showed fluctuating iIFAT titers, but demonstrated a consistent increase in VNA titers until termination of the experiment (1:12.7, 28 dpi). A similar pattern was observed in a surrogate assay mimicking virus neutralization (sVNT). Using the RBD of the SARS-CoV-2 spike-protein we further characterized antibody responses in an in-house ELISA (Figure 3, panels B-G, panel I). Anti-RBD IgM and IgG levels peaked at 8 and 12 dpi, respectively. A similar kinetics was observed in the infected contact animals #10 and #11. RBD-specific IgG2 patterns were highly similar to those of total IgG and total RBD antibodies. IgG2 antibodies with high neutralizing capacity had also been reported in dogs and their abundance correlates with neutralizing capacity *(26)* (Table 1, animals #5 and 6, 8 dpi). Although the amount of IgA in serum was limited (Figure 3, panel F), a similar trend as detected for the overall serum antibody levels was observed (Figure 3, panel G), e.g. with animal #6 having the highest values, and contact animals #10 and #11 reaching peak levels at later time points. RBD-specific antibodies were also detected in saliva samples 8 and 12 dpi (Figure 3, panel H-I) from animals that developed serum antibodies.

**Table 1.** Serological response of raccoon dogs to SARS-CoV-2 infection using the indirect immunoflourescence assay (iIFA), the virus neutralization test (VNT) and a surrogate Virus Neutralization test (sVNT). Positive results are highlighted in red (bold font) for inoculated (#1-9) and contact (#10-12) animals. No serological response on day 0 and day 4 pi (data not shown).

| No. | Day 8pi |  |  | Day 12pi |  |  | Day 16pi |  |  | Day 21pi |  |  | Day 28pi |  |  |
| --- | --- | --- | --- | --- | --- | --- | --- | --- | --- | --- | --- | --- | --- | --- | --- |
|  | iIFA | VNT | sVNT | iIFA | VNT | sVNT | iIFA | VNT | sVNT | iIFA | VNT | sVNT | iIFA | VNT | sVNT |
| #1 |  |  |  |  |  |  |  |  |  |  |  |  |  |  |  |
| #2 |  |  |  |  |  |  |  |  |  |  |  |  |  |  |  |
| #3 | **1:128** | < 1:4 | **56.39** |  |  |  |  |  |  |  |  |  |  |  |  |
| #4 | < 1:20 | < 1:2 | 13.42 |  |  |  |  |  |  |  |  |  |  |  |  |
| #5 | **1:64** | < 1:2 | **52.26** | **1:64** | < 1:2 | **71.62** |  |  |  |  |  |  |  |  |  |
| #6 | **1:128** | **1:5.04** | **52.19** | **1:64** | **1:12.7** | **83.99** |  |  |  |  |  |  |  |  |  |
| #7 | **1:128** | < 1:4 | **38.94** | **1:64** | < 1:2 | **72.56** | **1:64** | **1:4** | **76.22** | **1:128** | **1:10.08** | **84.99** | **1:1024** | **1:12.7** | **88.57** |
| #8 | < 1:20 | < 1:2 | 6.62 | < 1:20 | < 1:2 | 9.20 | < 1:20 | < 1:2 | 10.58 | < 1:20 | < 1:2 | 4.99 | < 1:20 | < 1:2 | -8.80 |
| #9 | < 1:20 | < 1:2 | 2.12 | < 1:20 | < 1:2 | 16.08 | < 1:20 | < 1:2 | 11.42 | < 1:20 | < 1:2 | 3.99 | < 1:20 | < 1:2 | -2.88 |
| #10 | < 1:20 | < 1:2 | 3.02 | < 1:20 | < 1:2 | **24.62** | **1:64** | < 1:2 | **62.34** | **1:128** | < 1:4 | **82.31** | **1:512** | < 1:4 | **82.98** |
| #11 | < 1:20 | < 1:2 | -9.55 | < 1:20 | < 1:2 | **47.88** | **1:64** | < 1:2 | **71.94** | **1:128** | **1:5.04** | **69.10** | **1:256** | < 1:4 | **81.82** |
| #12 | < 1:20 | < 1:2 | -0.77 | < 1:20 | < 1:2 | 8.89 | < 1:20 | < 1:2 | 12.17 | < 1:20 | < 1:2 | **22.91** | < 1:20 | < 1:2 | 7.34 |

**Figure 3.**
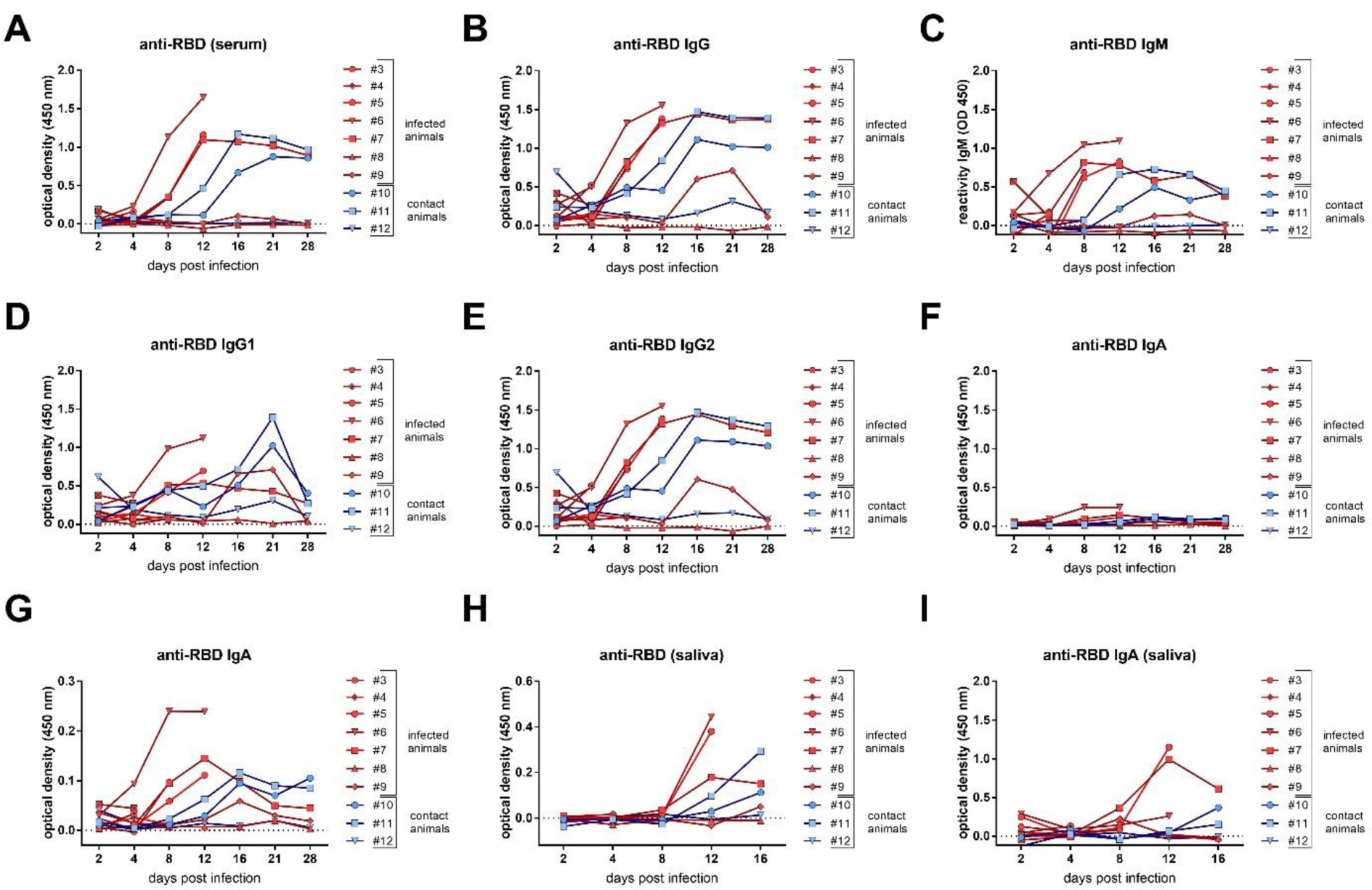
SARS-Co-V-2-specific antibody response (A) Individual immune response in sera as measured using an in-house RBD ELISA with a multi species conjugate (SBVMILK, kindly provided by ID-VET, Grabels, FRANCE); (B) with a secondary anti-dog IgG-conjugate; (C) with a secondary anti-dog IgM; (D) with a secondary anti-dog IgG1; (E) with a secondary anti-dog IgG2; (F) with RBD as antigen and a secondary anti-dog IgA, at the same scale, and (G) at a zoomed scale. (H) Total anti-RBD antibodies in saliva and (I) anti-RBD IgA in saliva

To test whether viral adaptions occurred during infection of raccoon dogs with this human SARS-CoV-2 isolate, we performed high throughput sequencing of SARS-CoV-2 re-isolated from nasal swabs of infected raccoon dog #2 at 2 dpi and contact animal #10 at 8 dpi yielding 100% identity to the inoculum (2019_nCoV Muc-IMB-1).

## Discussion

The present experimental study demonstrates that raccoon dogs are susceptible to SARS-CoV-2 infection and transmit the virus to contact animals. Six out of nine animals were successfully infected by intranasal inoculation. The susceptibility of raccoon dogs thus appears similar to Rousettus bats (*Rousettus aegyptiacus*) and slightly lower than ferrets (*Mustela putorius furo*) *(23) (23)*. Virus shedding in nasal and oropharyngeal swabs of raccoon dogs resulted in successful onward transmission of SARS-CoV-2 to two out of three contact animals as has been observed for other animal species with direct cage neighbors *(23, 29–31)*.

Increasing evidence supports the potential of several carnivore species to become infected by SARS-CoV-2 as a result of anthropo-zoonotic transmission *(13)*, possibly leading to re-infections of humans *(13)*. Therefore, wild carnivore species whether free-living or held in captivity should also be considered as intermediate hosts. With China’s substantial contribution to the global fur production of > 50 million animals per annum (0) (Appendix Figure3, panel A), it is conceivable that raccoon dogs may have played a hitherto unexplored role in the development of the pandemic, particularly considering the very mild signs of infection, efficient replication and transmission, and genetic stability. These environments with close contact between animals and an obvious interface with humans support SARS-CoV-2 transmission as was seen in several large mink farm outbreaks in The Netherlands, Denmark and Spain *(13, 19, 32, 33)*.

No obvious clinical signs could be observed, which is in line with experimental studies in other carnivores, i.e. adult cats (Felis catus) and ferrets that showed productive SARS-CoV-2 infection with no, or only mild clinical signs *(23, 31)*. By prominent nasal virus shedding in the absence of symptoms, raccoon dogs present a picture of infection resembling asymptomatic infections in other animals reflected by restriction of virus replication to the upper respiratory tract, substantiating the role of the nasal cavity in infection as shown for other animal species *(23, 31)*, as well as the majority of human cases *(34)*. Except for a mild rhinitis associated with the presence of viral antigen in the nasal mucosa, no other infection-related histopathological changes were observed. However, the absence of viral genome, pathohistological changes or viral antigen in the lungs of infected animals argue against raccoon dogs as a model for pulmonary manifestation of COVID-19.

The serological results suggest that the induction of SARS-CoV-2 specific VNA in raccoon dogs is reduced compared to ferrets but comparable to Egyptian fruit bats *(23)*. A delayed production of VNAs cannot be excluded, but appears unlikely against the dynamic increase of the measured ELISA antibodies. A mucosal immune response to SARS-CoV-2, i.e. antibodies in saliva were detected in raccoon dogs already 12 dpi (Figure 3), supporting the use of saliva as an early and non-invasive sample for epidemiological studies *(35)*. The limited presence of viral antigen in infected raccoon dogs at 4 dpi and the rapid decrease in viral loads prior to the development of measurable humoral immunity indicates that innate immune responses including interferon, mucus movement and epithelial cell turnover may play a prominent role in reducing infection.

High throughput sequencing of SARS-CoV-2 re-isolated from nasal swabs of infected raccoon dogs and contact animals yielded 100% identity to the inoculum (2019_nCoV Muc-IMB-1), demonstrating that no mutations occurred during virus replication in raccoon dogs. This is in contrast to findings in infected ferrets where two nonsynonymous single nucleotide exchanges after the ferret passage were identified *(25)*. This may indicate that the virus is already sufficiently adapted to this putative intermediate host.

In conclusion, further evidence is required from research about the origin of this pandemic. Large-scale sero-epidemiological studies in susceptible animals are needed. Historical samples collected prior to the epidemic are of particular importance and should preferentially also include a time series of archived samples. Further, affected fur farms may serve as reservoirs for SARS-CoV-2 and this risk should be mitigated by efficient and continuous surveillance. While SARS-CoV-2 might be controlled in holdings by very strict measures *(13, 32)*, a spill-over into susceptible wildlife species, in particular free living raccoon dogs representing one of the most successful invasive carnivore species in Europe *(6)* (Appendix Figure 3, panel B), would be even a greater challenge for elimination as long as preventive options are limited.

## Acknowledgments

We acknowledge Jeannette Kliemt, Mareen Lange, Silvia Schuparis, Gabriele Czerwinski, Bianka Hillmann and Patrick Zitzow for their technical assistance and Frank Klipp, Doreen Fiedler, Harald Manthei, René Siewert, Christian Lipinski, Ralf Henkel and Domenique Lux for their support during animal experiments. Funding: Intramural funding by the German Federal Ministry of Food and Agriculture was provided to the Friedrich-Loeffler-Institut. The funder of the study had no role in study design, data collection, data analysis, data interpretation, or writing of the report. T.C.M and M.B had full access to all the data in the study and had final responsibility for the decision to submit for publication.

## Disclaimers

The authors declare no competing interests.

