## Supplement for "Susceptibility of raccoon dogs for experimental SARS-CoV-2 infection"

### Appendix

#### Material and Methods

Animals & study design

The experimental design followed the general setup of a previous study (*1)*. Fourteen adult, male (n=4) and female (n=10) raccoon dogs originating from a commercial farm were used. All animals were tested negative by RT-qPCR and antibody tests (ELISA, indirect immunofluorescence assay iIFAT, virus neutralization test VNT) for SARS-CoV-2 prior to the experiment.

All raccoon dogs had been vaccinated against distemper, adeno- and parvovirus (Eurican® SHP, Merial, France). Animals were kept in individual stainless-steel cages (1.5m x 0.95m x 2.0m) in four separate segments at 20°C room temperature, 60-80% humidity and a 12hr/12hr (35% dimming during night modus) lighting control within a fan forced draught ventilation equipped BSL3** animal facility at the Friedrich-Loeffler-Institut (FLI). Water was offered ad libitum. Animals were fed daily with 400 gr commercially produced feed for farmed foxes and raccoon dogs (Schirmer und Partner GmbH Co KG, Germany; Michael Hassel GmbH, Langenargen, Germany). The diet was supplemented with vitamins, minerals and items like one-day old chickens (*2)* The general health status of all animals, feed uptake and defecation were recorded daily. The body weight and temperature of all animals were measured prior to inoculation and at days 2, 4, 8, 12, 16, 21, and 28 pi.

Nine raccoon dogs (3 males, 6 females) were infected intranasally with $10^5$ $TCID_{50}$ SARS-CoV-2 2019_nCoV Muc-IMB-1. The inoculum of 2x1ml was administered to both nostrils using a pipette. To test viral transmission by direct contact, three naïve sentinel animals (all female) were added 24 hours post inoculation. Nasal, oropharyngeal and rectal swabs were taken at 2, 4, 8, 12, 16, 21 and 28 days post infection (dpi), blood was taken at 4, 8, 12, 16, 21 and 28 dpi. Two animals each were sacrificed at day 0 (control #1, #2) day 4 (animals #1, #2), day 8 (animals #3, #4) and day 12 pi (animals #5, #6). The remaining inoculated animals (animals #7-9) and the contacts (animals #10-12) were euthanized 28 dpi. All animals were subjected to autopsy for macroscopic evaluation and tissue sampling. The animal experiments were evaluated and approved by the ethics committee of the State Office of Agriculture, Food Safety, and Fishery in Mecklenburg – Western Pomerania (LALLF M-V: LVL MV/TSD/7221.3-2-010/18-12).

Virus and cells

SARS-CoV-2 isolate 2019_nCoV Muc-IMB-1 was kindly provided by R. Woelfel (German Armed Forces Institute of Microbiology, Munich, Germany). The complete sequence of this isolate is available through GISAID under the accession ID_EPI_ISL_406862 and designation “hCoV-19/Germany/BavPat1/2020”. The virus was propagated once in Vero E6 cells in a mixture of equal volumes of Eagle MEM (Hanks’ balanced salts solution) and Eagle MEM

(Earle's balanced salts solution) supplemented with 2mM L-Glutamine, nonessential amino acids, adjusted to 850 mg/L, NaHCO3, 120 mg/L sodium pyruvate, 10% fetal bovine serum (FBS), pH 7.2. No contaminants were detected within the virus stock preparation by metagenomic analysis employing previously published high throughput sequencing procedures Ion Torrent S5XL instrument (*3, 4)* and the sequence identity of the passaged virus (study accession number: PRJEB39640) was confirmed. The virus was harvested after 72 h, titrated on Vero E6 cells and stored at -80°C until further use.

#### RNA extraction and detection of SARS-CoV-2

Total RNA from nasal, oropharyngeal and rectal swab samples, fecal samples as well as from tissues taken at autopsy were extracted using the NucleoMagVet kit (Macherey&Nagel, Düren, Germany) according to manufacturer's instructions. Tissues were homogenized in 1 ml cell culture medium (see above) and a 5 mm steel bead in a TissueLyser (Qiagen, Hilden, Germany). Fecal samples were vortexed in sterile NaCl and the supernatant was sterile filtered (22µm) after centrifugation. Swab samples were transferred into 1 ml of serum-free tissue culture media and further processed after 30 min shaking.

SARS-CoV-2 RNA was detected by an E-gene based RT-qPCR (*5).* The RT-qPCR reaction was prepared using the AgPath-ID-One-Step RT-PCR kit (Thermo Fisher Scientific, Waltham, Massachusetts, USA) in a volume of 12.5 µl including 1 µl of ß-Actin-mix2-HEX as internal control and 2.5 µl of extracted RNA. The reaction was performed for 10 min at 45°C for reverse transcription, 5 min at 95°C for activation, and 42 cycles of 15 sec at 95°C for denaturation, 20 sec at 57°C for annealing and 30 sec at 72°C for elongation. Fluorescence was measured during the annealing phase. All RT-qPCRs were performed on a BioRad real-time CFX96 detection system (Bio-Rad, Hercules, USA). Absolute quantification was done using a standard quantified by the QX200 Droplet Digital PCR System in combination with the 1-Step RT-ddPCR Advanced Kit for Probes (BioRad, Hercules, USA).

Nasal swab samples from raccoon dog #2 (2 dpi) and from contact animal #10 (8 dpi) were subjected to high-throughput sequencing and compared to the inoculum (study accession number: PRJEB39640) by employing previously published high throughput sequencing procedures using Ion Torrent S5XL instrument (*3, 4).*

#### Detection of SARS-CoV-2 reactive antibodies

For iIFAT, confluent Vero E6 cells in a 96 well plate were infected with 0.1 MOI of SARS-CoV-2 or overlaid with cell culture medium for negative control cells. After 24h, cells were fixed with 4% paraformaldehyde and permeabilized with 0.5% Triton-X-100. Serum samples were heat inactivated at 56°C for 30 min. For antibody detection, 50 µl of a 2-fold dilution series of the serum samples (starting from 1:20) were added in parallel to the SARS-CoV-2 positive and negative cells. After 1h incubation, cells were washed and incubated for 1h with a goat-α-dog-IgG-FITC antibody (1:250, Bethyl, Texas, USA). After final washing, cells were analyzed by fluorescence microscopy.

For VNT, 50 µl of medium containing $10^{3.3}$ $TCID_{50}$ SARS-CoV-2 were mixed with 50 µl of serially diluted serum. Each sample was tested in triplicates. After 1h incubation at 37°C the mixture was transferred to confluent Vero E6 cells in a 96 well plate. Viral replication was assessed after 5 days at 37°C, 5% $CO_2$ by the detection of CPE.

For ELISA, the SARS-CoV2 RBD-SD1 domain (amino acids 319 – 519 of the SARS-CoV2 Spike ectodomain) was amplified from a codon-optimized synthetic gene (GeneArt, Thermo Scientific). The construct was cloned into the expression vector pEXPR103 (iba lifesciences) in frame with an N-terminal modified mouse Ig kappa light chain signal peptide and a C-terminal double Strep tag. Expi293 cells were grown in Expi293 expression medium (Thermo Scientific) and polycarbonate Erlenmeyer flasks (Corning) at 37°C, 8% $CO_2$, 125 rpm. For transfection, cell density was adjusted to 2 x $10^6$ cells/ml and a total volume of 70 ml was transfected with 70 µg of plasmid DNA using the ExpiFectamine293 transfection kit (Thermo Scientific) according to the manufacturer´s instructions. The cells were subsequently incubated at 37°C, 8% $CO_2$, 125 rpm. The supernatant was harvested five days after transfection by centrifugation at 6000 x g for 20 min at 4 °C. Biotin was blocked by addition of BioLock (iba lifesciences) as recommended, and the supernatant was purified using Strep-Tactin XT Superflow high capacity resin (iba lifesciences) according to the protocol of the manufacturer. The proteins were eluted with 50 mM Biotin (in 100 mM Tris-HCl, 150 mM NaCl, 1mM EDTA; pH 8.0) and stored at -80 °C until further use. Medium-binding ELISA plates (Greiner Bio-One GmbH, Germany) were coated with 100 ng/well of the SARS-CoV-2 RBD overnight at 4 °C in 0.1 M carbonate buffer at pH 9.6 or treated with the coating buffer only. Thereafter, the plates were washed three times using Tris-buffered saline with Tween (TBST) and blocked for 1 h at 37 °C using 5% skimmed milk in phosphate-buffered saline (PBS). The sera were diluted 1:100 in TBST and incubated on the coated and uncoated wells for 1h at room temperature followed by three washes using TBST. The saliva samples were used undiluted. Reactivity was shown by adding a multi species conjugate (SBVMILK; IDvet, France) diluted 1:80 (serum) or 1:10 (saliva). After an incubation period of 1 h at room temperature, the plates were washed again and Tetramethylbenzidine (TMB) substrate (IDEXX, Switzerland) was added. The ELISA readings were taken at a wavelength of 450 nm on a Tecan Spectra Mini instrument (Tecan Group Ltd, Switzerland). The measurements were normalized to the respective samples tested on wells treated only with the coating buffer.

For comparison, sera were also tested in a newly developed commercial SARS-CoV-2 sVNT designed to detect total neutralizing antibodies in an isotype- and species-independent manner (GenScript USA). The test is based on antibody-mediated blockage of virus-host interaction between the ACE2 receptor protein and the RBD of the viral S protein (*6)*. Briefly, 1:10 serum dilutions were incubated for 30 min at 37°C with HRP-coupled RBD before transferring the samples to the capture plate pre-coated with the human ACE2 protein. After 15 min incubation at 37°C, plates were washed four times. TMB substrate was added and the plate was incubated at room temperature for 15 min before stopping the reaction and reading the optical density (OD) at 450 nm. Percent inhibition was calculated as (1- OD sample / OD negative control) x 100.

Pathology: Autopsy, histopathology, immunohistochemistry

The following tissues were collected and fixed in 10% neutral-buffered formalin and trimmed for paraffin embedding: nasal atrium (non-respiratory region), nasal conchae (respiratory and olfactory region, cross sections approximately every 5 mm after decalcification), soft palate, tonsil, parotid and mandibular salivary gland, trachea (upper and lower third), lung (inflated with formalin, cross sections of the right cranial, medial, caudal and accessory lobe), tracheobronchial lymph node, heart, liver with gall bladder, spleen, stomach (cardiac, fundic, pyloric region), intestine (duodenum, jejunum, ileum, colon, cecum, rectum), pancreas, kidney, ureter, urinary bladder, adrenal gland, skeletal muscle, skin, and brain (3 coronal section with lobus olfactorius, hippocampus and cortex, cerebellum und medulla oblongata). In addition, blood (5 ml EDTA, rest serum), urine (puncture at autopsy), cerebrospinal fluid, and bronchio-alveolar lavage (left lobe, using 10 ml PBS), were taken for viral RNA detection and virus isolation.

The nasal mucosa was particularly scored for cellular degeneration and necrosis, loss of epithelium, intraluminal debris, mucosal infiltration and transmigration, endothelial activation (swelling), edema, hemorrhage, re-epithelization/squamous metaplasia and granulation tissue formation. All lung sections were scored for lesion on the pleural surface, alveolar collapse, endothelial activation (swelling), edema (interstitial, bronchial, peribronchial, perivascular, alveolar), congestion, inflammatory infiltrates (interstitial, peribronchial, perivascular), increased numbers of alveolar macrophages, necrosis (bronchial, alveolar), squamous metaplasia, hyperplasia (bronchial, pneumocyte type 2), presence of multinucleated giant cells, cellular debris in bronchial lumina, alveolar emphysema.

### Statistical information

No statistical methods were used. The experiments were not randomized and the investigators were not blinded to allocation during experiments and outcome assessment. All samples for downstream virological and serological analyses were blinded. All data were analyzed and visualized using GraphPad Prism Version 7.0 (GraphPad Software, San Diego, CA, USA).

6. Tan CW, Chia WN, Chen MI-C, Hu Z, Young BE, Tan Y-J, et al. *A SARS-CoV-2 surrogate virus neutralization test (sVNT) based on antibody-mediated blockage of ACE2-spike (RBD) protein-protein interaction*. Epub ahead of print. doi: 10.21203/rs.3.rs-24574/v1.

### Supplemental Figures

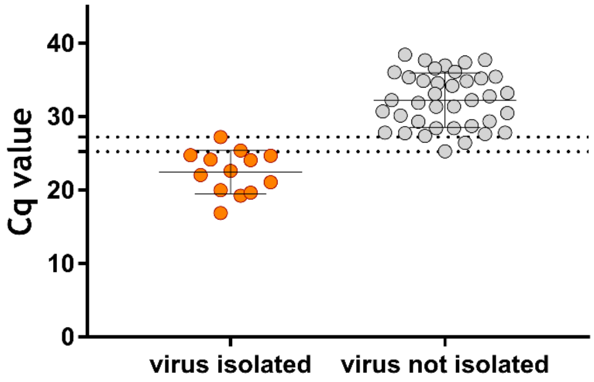


**Appendix Figure 1. Graphical display of Cq values of swabs, stratified by successful virus isolation.** Dashed lines represent the upper bound of cq-values for successful virus isolation and the lower bound of cq-values for which virus isolation was not successful

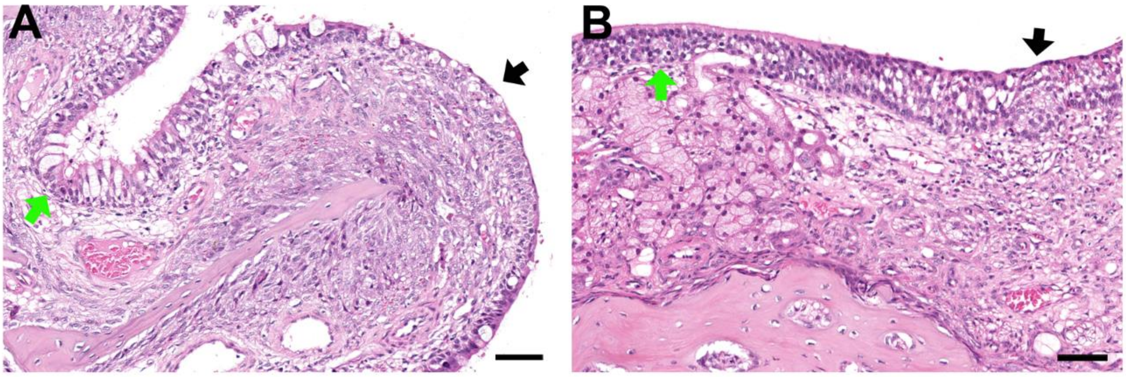


**Appendix Figure 1. Lesions in the nasal mucosa, 28 days after experimental SARS-CoV-2 infection of racoon dogs**. (**A**) Racoon dog #7 at 28 dpi, respiratory region, unaffected respiratory epithelium (green arrow) and affected area with squamous metaplasia and granulation tissue formation (black arrow), (**B**) Raccoon dog #7 at 28 dpi, olfactory region, unaffected olfactory epithelium and subepithelial Bowman's glands (green arrow) and affected irregular olfactory epithelium with loss of Bowmans's glands and granulation tissue formation. Histopathology, hematoxylin & eosin stain, all bar 50 µm. Note: lesions were detected in successfully infected animals only (#7, 11), but viral antigen detection by immunohistochemistry failed 28 dpi. Thus, findings should be interpreted with caution and are only indication for previous replication sites.

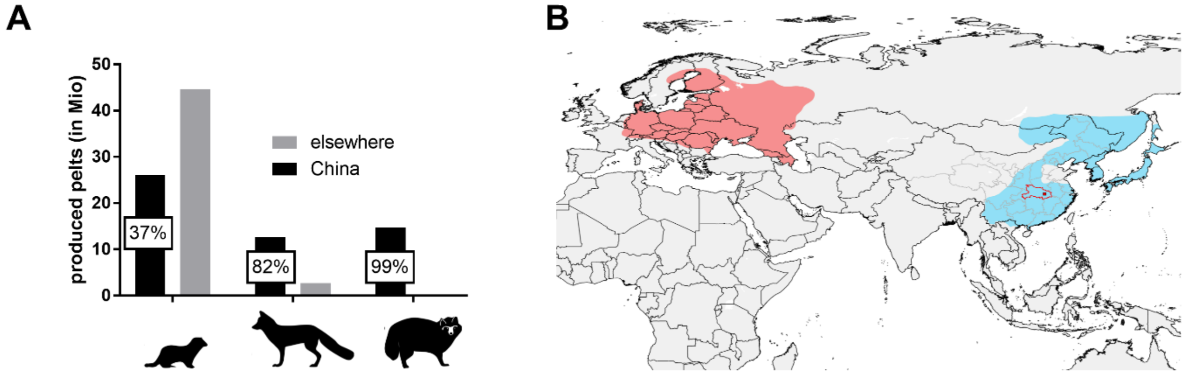


**Appendix Figure 3. Relevance of fur production, worldwide distribution of racoon dogs and phylogenetic overview of potential animal hosts for SARS-CoV-2**. (A) Total number of produced pelts of minks, foxes and raccoon dogs in China and elsewhere in the world. The percentage of China's share is indicated. Data source: China's fur trade and its position in the global fur industry, ACTAsia 2019. (B) Traditional distribution (blue) and anthropogenic expansion (orange) of the raccoon dog (*Nyctereutes procyonoides*, Data source: IUCN list of species, 2016. Nyctereutes procyonoides. https://www.iucnredlist.org/species/14925/85658776). The location of the Hubai district in China is indicated in red.

**Supplementary pathology findings interpreted as non-related to SARS-CoV-2 infection**

At autopsy, several animals exhibited oligofocal pulmonary atelectasis. Histopathology identified alveolar collapse without significant inflammatory infiltrates in affected regions, but also in otherwise macroscopically inconspicuous lungs. The histologic alteration was also found in negative control animals as well as in animals inoculated with SARS-CoV-2 that were tested negative for the virus and seroconversion throughout the experiment. Viral antigen could not be detected at any time point in any lung tissue tested. Thus, this finding was interpreted as not-related to SARS-CoV-2 infection.

Animal #6, showed rare necrosis of bronchial and alveolar epithelium, cellular debris in bronchial lumina, multifocal to coalescing hyperplasia as well as squamous metaplasia of the bronchial epithelium, and multifocal to diffuse peribronchial edema. Inflammation in the lung but also in the gall bladder was characterized by diffuse, mainly lymphocytic and eosinophilic inflammatory infiltrates. The right, medial lobe of the liver of this animals was affected by cholangiocellular carcinoma. Taken together, the findings in the lung, gall bladder and liver are interpreted to be related to a chronic parasitic burden, most likely trematodes.

Additional findings, in the lung and all other tissues tested did not exceed lesions found in the negative control animals.
